# Genomic data enables genetic evaluation using data recorded on low-middle income country smallholder dairy farms

**DOI:** 10.1101/827956

**Authors:** Owen Powell, Raphael Mrode, R. Chris Gaynor, Martin Johnsson, Gregor Gorjanc, John M. Hickey

**Affiliations:** The Roslin Institute and Royal (Dick) School of Veterinary Studies, University of Edinburgh, Easter Bush, Midlothian, EH25 9RG, UK; Scotland’s Rural College (SRUC), Easter Bush, Midlothian, EH25 9RG, UK; International Livestock Research Institute (ILRI), Nairobi 00100, Kenya; Department of Animal Breeding and Genetics, Swedish University of Agricultural Sciences, Box 7023, 750 07, Uppsala, Sweden

## Abstract

**Background:** Genetic evaluation is a central component of a breeding program. In advanced economies, most genetic evaluations depend on large quantities of data that are recorded on commercial farms. Large herd sizes and widespread use of artificial insemination create strong genetic connectedness that enables the genetic and environmental effects of an individual animal’s phenotype to be accurately separated. In contrast to this, herds are neither large nor have strong genetic connectedness in smallholder dairy production systems of many low to middle-income countries (LMIC). This limits genetic evaluation, and furthermore, the pedigree information needed for traditional genetic evaluation is typically unavailable. Genomic information keeps track of shared haplotypes rather than shared relatives. This information could capture and strengthen genetic connectedness between herds and through this may enable genetic evaluations for LMIC smallholder dairy farms. The objective of this study was to use simulation to quantify the power of genomic information to enable genetic evaluation under such conditions.

**Results:** The results from this study show: (i) the genetic evaluation of phenotyped cows using genomic information had higher accuracy compared to pedigree information across all breeding designs; (ii) the genetic evaluation of phenotyped cows with genomic information and modelling herd as a random effect had higher or equal accuracy compared to modelling herd as a fixed effect; (iii) the genetic evaluation of phenotyped cows from breeding designs with strong genetic connectedness had higher accuracy compared to breeding designs with weaker genetic connectedness; (iv) genomic prediction of young bulls was possible using marker estimates from the genetic evaluations of their phenotyped dams. For example, the accuracy of genomic prediction of young bulls from an average herd size of 1 (μ=1.58) was 0.40 under a breeding design with 1,000 sires mated per generation and a training set of 8,000 phenotyped and genotyped cows.

**Conclusions:** This study demonstrates the potential of genomic information to be an enabling technology in LMIC smallholder dairy production systems by facilitating genetic evaluations with *in-situ* records collected from farms with herd sizes of four cows or less. Across a range of breeding designs, genomic data enabled accurate genetic evaluation of phenotyped cows and genomic prediction of young bulls using data sets that contained small herds with weak genetic connections. The use of smallholder dairy data in genetic evaluations would enable the establishment of breeding programs to improve *in-situ* germplasm and, if required, would enable the importation of the most suitable external germplasm. This could be individually tailored for each target environment. Together this would increase the productivity, profitability and sustainability of LMIC smallholder dairy production systems. However, data collection, including genomic data, is expensive and business models will need to be carefully constructed so that the costs are sustainably offset.

## Background

The huge increase in milk yield of dairy cattle in advanced economies over the past century is a powerful example of the impact that selective breeding can have on improving livestock productivity. For example, in the US dairy industry, production of milk per cow doubled from an average of 20 litres to 40 litres per day between 1960 and 2000 [1]. Approximately 50% of this improvement can be attributed to breeding. However, despite the potential benefits, similar breeding practices have had poor efficacy and adoption in smallholder dairy production systems in many low to middle-income countries (LMICs). Recent estimates from Kenyan smallholder farms suggest that average productivity per cow is approximately 5 litres per day and there is little evidence of major genetic improvement in recent decades [2–5].

In Kenya and other East African countries, farms with five cows or less account for more than 70% of milk production [6,7], and farms with 10 cows or less account for around 90% of milk production [8]. The low levels of productivity and its economic importance has stimulated renewed efforts to improve dairy cow productivity in LMIC smallholder dairy production systems [6,9–11]. These efforts include new approaches for collecting data from rural farms more effectively and the establishment of effective and penetrant genetic evaluation schemes [10,12–14], breeding programs and dissemination programs [15], all of which have been somewhat intractable to sustain over the long-term in the past.

Genetic evaluation is a central component of a breeding program. The properties of an ideal data set that enables an accurate genetic evaluation include: (i) genetic connectedness between herds or management groups [16]; (ii) sufficient numbers of animals; (iii) sufficiently large herd sizes; and (iv) accurate phenotype collection. Genetic evaluations have been very successful in advanced economies because large data sets are routinely assembled from commercial farms with modest to large herd sizes (e.g., twenty to several thousand cows). Genetic connectedness between herds is high due to the widespread use of artificial insemination (AI). Typically, phenotypes are accurately measured (e.g., automatically on advanced milking machines). Such data enables the genetic and environmental effects of an individual animal’s phenotype to be accurately separated. All or many of these features are not present in many LMIC smallholder dairy production systems. For example, smallholder dairy farmers in East Africa have small herd sizes (e.g., herds with one to five cows), a low prevalence of AI (5-10%) [8], and an absence of automated phenotyping systems [17]. Traditionally, this has prevented the establishment of effective genetic evaluation systems in these settings.

Genomic evaluations use a genomic relationship matrix to capture the realised, rather than expected pedigree-derived relationships between animals [18,19]. The use of genomic information has been transformative for many genetic evaluation systems in advanced economies. For example, the accuracy, which is the square root of reliability, of prediction for milk yield of young bulls increased from 0.62 using pedigree best linear unbiased prediction (PBLUP) to 0.85 for genomic best linear unbiased prediction (GBLUP) [20]. In the context of LMIC smallholder dairy production systems, genomic data could be even more important than it has been in advanced economies. For the first time, genomic data could enable effective genetic evaluation systems based on relatively imprecisely measured phenotypes, collected on cows in very small herd sizes, which have relatively low levels of genetic connectedness. In such a setting, genomic data could capture and utilise information pertaining to haplotypes that are shared by animals in different herds. This information could reveal genetic connectedness that is unseen by pedigree information, which would, in turn, enable more accurate partitioning of the genetic and environmental effects on animal’s performance in small herds. This opens up the possibility of an *in-situ* breeding program based on *in-situ* performance data from LMIC smallholder dairy farms. Given that such data reflects the performance of animals within the target management and environment settings, animals produced by such a breeding program would be most suited to the participating smallholder dairy farmers.

In genetic evaluations, the herd or management group is usually included in the statistical model to enhance the separation of the genetic and environmental effects of an animal’s performance [21–24]. Herds can be modelled as fixed or random effects. Most genetic evaluations in advanced economies model herds as fixed effects because herd sizes are typically large, which leads to fixed and random effects models giving almost equal solutions [22,23]. When herd sizes are small, such as in many LMIC smallholder dairy production systems, modelling herd as a fixed effect leads to inaccurate solutions [25]. Modelling small herds as random effects may reduce this inaccuracy, providing estimated breeding values (EBVs) with higher accuracies. In combination with the use of genomic information, this could enable genetic evaluations to be performed using data recorded, *in-situ*, on LMIC smallholder dairy farms.

The aims of this study were to use simulation to quantify: (i) the power of genomic information to enable genetic evaluation based on phenotypes recorded on smallholder dairy farms and, under such conditions, the impact of: (ii) modelling herd as a fixed or random effect; (iii) the genetic connectedness of a breeding population; and (iv) the number of records on the accuracy of EBVs of phenotyped cows and young bulls.

Across a range of breeding designs, genomic data enabled accurate genetic evaluation of phenotyped cows using data sets that contained small herds with weak genetic connections (according to pedigree). The genetic evaluation of phenotyped cows using genomic information had higher accuracy compared to pedigree information across all breeding designs. The genetic evaluation of phenotyped cows with genomic information and modelling herd as a random effect had higher or equal accuracy compared to modelling herd as a fixed effect. The genetic evaluation of phenotyped cows from breeding designs with strong genetic connectedness had higher accuracy compared to breeding designs with weaker genetic connectedness. The genomic prediction of young bulls was possible using marker estimates from the genetic evaluations of their phenotyped dams. For example, the accuracy of genomic prediction of young bulls from an average herd size of 1 (μ=1.58) was 0.40 under a breeding design with 1,000 sires mated per generation and a training set of 8,000 phenotyped and genotyped cows. Our results show that genetic evaluations with genomic information can provide a high accuracy of EBVs of phenotyped cows and young bulls when using data from smallholder dairy farms, and would, therefore, enable *in-situ* breeding programs based on performance measured *in-situ*.

## Material and methods

Simulations were used to quantify the power of genomic information to enable genetic evaluation based on phenotypes recorded on smallholder dairy farms. Ten replicates of several scenarios were performed with the overall simulation scheme depicted in Figure 1. The simulations were performed using AlphaSimR [26] and were designed to: (i) generate whole genome sequence data; (ii) generate single nucleotide polymorphisms (SNP), quantitative trait loci (QTL) and phenotypes; (iii) generate pedigree structures for LMIC smallholder dairy populations; (iv) vary the population and average herd size; (v) vary the ratios of genetic, herd and environmental variances; and (vi) run genetic evaluations modelling herd as either fixed or random effects. Conceptually, the simulation scheme was divided into historical and evaluation phases.

**Figure 1.**
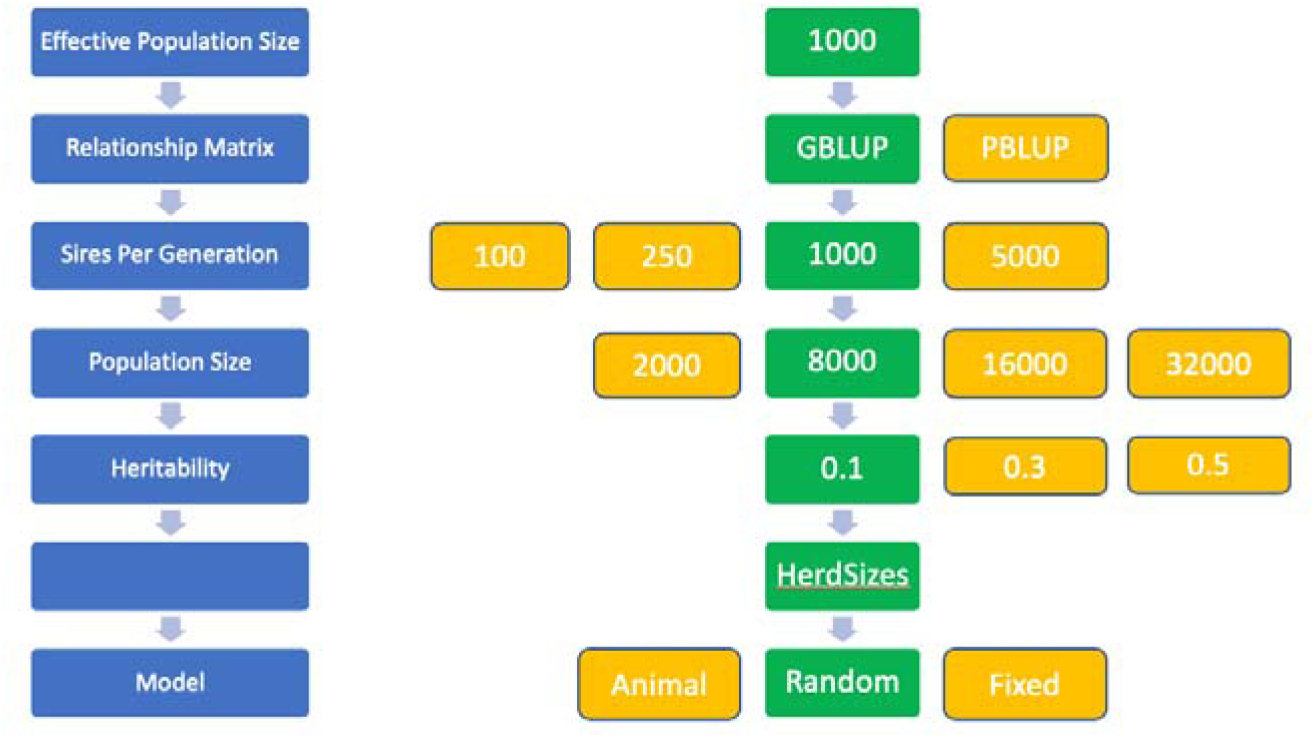
Simulation scenarios. The conventional breeding design is highlighted in green. Breeding designs were compared for each design parameter individually (horizontally), while keeping all other design parameters fixed at the values of the conventional breeding design.

Each of the 10 replicates consisted of: (i) a burn-in phase shared by all strategies; and (ii) an evaluation phase that simulated breeding with each of a number of different breeding designs. Specifically, the historical component was subdivided into three stages: the first simulated the species’ genome sequence; the second simulated founder genotypes for the initial parents; and the third simulated five generations of breeding using phenotypic selection.

The burn-in phase represented historical evolution, under the assumption that livestock populations have been evolving for tens of thousands of years, and historical breeding efforts that were represented by five generations of phenotypic selection. The evaluation phase represented six generations of animal breeding in which animals were selected on their phenotypes. In the evaluation phase, population parameters were varied (i.e., the number of sires mated per generation, large or small population sizes, large or small average herd sizes, and different proportions of the genetic, herd and environmental variances) to resemble a range of possible breeding designs (Figure 1).

### Burn-In: Generation of whole genome sequence data

For each replicate, a genome consisting of 10 chromosome pairs was simulated for the hypothetical animal species similar to cattle. Sequence data was generated using the Markovian Coalescent Simulator (MaCS) [27] and AlphaSimR [26] for 4,000 base haplotypes for each of ten chromosomes. The chromosomes were each 100 cM in length comprising 10^8^ base pairs and were simulated using a per site mutation rate of 1×10^-8^ and a per site recombination rate of 1×10^-8^. The N_e_ was set to 1,035 in the final generation of historical simulation, to N_e_=6,000 (1,000 years ago) to N_e_=24,000 (10,000 years ago), and to N_e_=48,000 (100,000 years ago) with linear changes in between [28]. The N_e_ of 1,035 was chosen to reflect the high genetic diversity found in cattle populations in Africa.

### Burn-In: Founder Genotypes

Simulated genome sequences were used to produce 2,000 founder animals. These founder animals served as the initial parents in the burn-in phase. Sites segregating in the founders’ sequences were randomly selected to serve as 5,000 SNP markers per chromosome (50,000 genome-wide in total) and 1,000 QTL per chromosome (10,000 genome-wide in total).

### Burn-In: Phenotype

A single trait representing total milk yield for a single lactation was simulated for all animals. The true breeding values (TBVs) were calculated by summing the average effects of the animal’s genotype at each QTL. QTL additive effects were sampled from a standard normal distribution, N(0,1), and linearly scaled to produce TBVs in the founder population with a variance 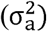 of 0.2. Random error was sampled from a normal distribution, 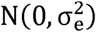. The initial random error variance was set at 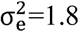. The TBVs and random error effects were summed to create the phenotypes of the animal. These phenotypes were used for selection during the burn-in and the first 5 years in the evaluation phases of the simulation. Additional herd effects were added to the phenotypes of the animals, described in a later section, in the final generation of the evaluation phase of the simulation

### Recent (Burn-In) Breeding

Recent (burn-in) breeding for milk yield was simulated over 5 discrete generations of selective breeding on phenotype. The features of this breeding stage were: (i) 225 sires per generation, (ii) 1,000 dams per generation, and (iii) 2,000 offspring per generation. These numbers were chosen to match the base population N_e_ of 1,035 following the equation from Charlesworth et al. (2008) that accounts for the variable number of males and females as well as the mean and variance of family size. In the final generation of this stage, 80,000 offspring were generated to enable the full range of scenarios in the evaluation phase of the simulation.

### Evaluation Phase

The evaluation phase of the simulation modelled breeding using alternative breeding designs. Each design was simulated for an additional 6 generations following the recent breeding burn-in component so that each design could be evaluated with an equivalent starting point. A baseline design was constructed using parameters that are representative of the current smallholder farming system commonly observed in East Africa. We refer to this design as the LMIC design. Alternative breeding designs were modifications that used the LMIC design as a template (Figure 1). The common features across the simulation of all the breeding designs were: (i) all generations of selection produced 80,000 animals of equal sex ratio, (ii) for simplicity selection on sires was based on their phenotype, (iii) no selection was performed on dams. Alternate breeding designs varied: (i) the size of the training set; (ii) the number of sires mated per generation; (iii) the average herd size; and (iv) the proportions of genetic, herd and environmental variances. A schematic for the overall structure of the breeding designs, including the LMIC design, is given in Figure 1 and a detailed description follows.

### LMIC Design

The LMIC design was developed to approximate the current smallholder farming system structure commonly observed in East Africa. The training set size was set at 8,000 phenotyped cows and the number of sires mated per generation was set to 1,000. A trait heritability of 0.1 and ratio of 1:4 between genetic and herd effect variance ratios were chosen based upon unpublished data [29].

### Genetic Evaluation Models

Breeding values were estimated using the following basic model:

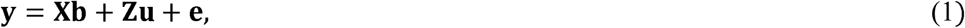

where **y** is a vector of phenotype records measured on cows; **b** is a vector of fixed effects; **u** is a vector of breeding values for which we assumed that with the PBLUP 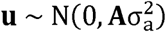 and with the GBUP 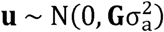, where **A** is the pedigree numerator relationship matrix based on 5 generations of the pedigree [30] and **G** is the genomic numerator relationship matrix based on 50k SNP chip [31]; **e** is a vector of residuals for which we assumes 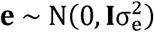; **X** and **Z** are the incidence matrices linking phenotype records respectively to **b** and **u**. We have conducted three analyses with the basic model in relation to a herd effect: (i) we excluded it, which gave us the basic model with intercept as the only fixed effect; (ii) we modelled it as a fixed effect; and (iii) we modelled it as a random effect for which we assumed 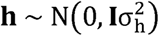. We assumed that the variance of herd effects 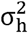, breeding values 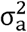 and residuals 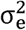 were known and set them to the simulated values of the LMIC design. Only the last generation of phenotype data was used in model 1 to mimic the recent introduction of phenotype, pedigree and genomic data recording.

PBLUP evaluations were run using the WOMBAT software [32]. GBLUP evaluations were run using the AlphaBayes software [33]. Three genetic evaluation models were fit: (i) excluding herd effects; (ii) herds modelled as fixed effects; and (iii) herds modelled as random effects. All models modelled the animal IDs as random effects. All other parameters were held constant at the values used in the LMIC design.

### Genetic Connectedness and Herd Size

Genetic connectedness was varied across different breeding designs in two ways; (i) herd connectivity – the distribution of related animals within and across different herds, and (ii) the recent N_e_ of the breeding design. The herd connectivity was varied by simulating different average herd sizes. To generate datasets with a range of different average herd sizes, the realised herd sizes were sampled from a Poisson distribution with a lambda of 1 (μ = 1.58, σ^2^ = 0.66), 2 (μ = 2.32, σ^2^ = 1.60), 4 (μ = 4.06, σ^2^ = 3.78), 8 (μ = 8, σ^2^ = 8), 16 (μ = 16, σ^2^ = 16.19) and 32 (μ = 32, σ^2^ = 31.92). The recent N_e_ of the breeding design was varied using four different numbers of sires mated per generation: 100, 250, 1,000 and 5,000 sires. The number of dams per generation remained constant at 40,000. All other parameters were held constant at the values used in the LMIC design.

### Size of Training Set

The size of the training set used in the genetic evaluations was varied across different breeding designs using four different numbers of records: 2,000, 8,000, 16,000 and 32,000 phenotyped cows. Phenotyped cows were sampled evenly across the population, to ensure the genetic connectedness was maintained. All other parameters were held constant at the values used in the LMIC design.

### Trait Heritability and Herd Effect

To produce the final phenotype records, the TBVs were standardized and re-scaled, and herd and random error effects were sampled from a normal distribution with corresponding variances. In addition to the LMIC design, which had a trait with a narrow sense heritability of 0.1 and herd effect variance ratio of 0.4, we simulated two other scenarios: (i) a trait with a narrow sense heritability of 0.3 and herd effect variance ratio of 0.4; and (ii) a trait with a narrow sense heritability of 0.5 and herd effect variance ratio of 0.4. For each of the three scenarios, the TBVs, herd effects and random errors were summed to create the final phenotypes of the cows. All other parameters were held constant at the values used in the LMIC design.

### Generation of young bull population

For each scenario we generated an additional generation of offspring to produce a validation set of 2,000, 8,000, 16,000 and 32,000 selection candidates, the young bulls that would have been genomically tested. Young bulls had no phenotypes recorded and as such served as forward validation of the model 1 fitted on phenotyped cows.

### Comparison of Breeding Designs

The various breeding designs resulted in 288 different scenarios which enabled multiple comparisons. The breeding designs were compared based upon the accuracy and bias of EBVs separately for each scenario and replicate – we report mean and 95% interval of estimates over replicates. Accuracy was measured as the Pearson’s correlation coefficient between the EBVs and TBVs. The bias of genomic prediction was measured as the slope of the regression of the TBVs on the EBVs.

## Results

The various breeding designs resulted in 288 different scenarios which enabled multiple comparisons. Across a range of breeding designs, genomic data enabled accurate genetic evaluation of phenotyped cows using data sets that contained small herds with weak genetic connections. The main trends observed in our results show: (i) the genetic evaluation of phenotyped cows using genomic information had higher accuracy compared to pedigree information across all breeding designs; (ii) the genetic evaluation of phenotyped cows with genomic information and modelling herd as a random effect had higher or equal accuracy compared to modelling herd as a fixed effect; (iii) the genetic evaluation of phenotyped cows from breeding designs with strong genetic connectedness had higher accuracy compared to breeding designs with weaker genetic connectedness; (iv) the genomic prediction of young bulls was possible using marker estimates from the genetic evaluations of their phenotyped dams. For example, the accuracy of young bulls from an average herd size of 1 (μ=1.58) was 0.40 under a breeding design with 1,000 sires mated per generation and a training set of 8,000 phenotyped and genotyped cows. The accuracies of genomic prediction of young bulls followed similar trends to those observed in the evaluation of phenotyped cows, with a reduction of ∼0.1 in overall accuracy.

To ease the presentation, we break the results into 5 sections: (i) LMIC design; (ii) impact of herd effect modelling; (iii) impact of genetic connectedness and heritability; (iv) impact of training set size; and (v) prediction of young bulls.

### LMIC Design

The accuracy of genetic evaluation of phenotyped cows, from small, weakly genetically connected herds was quantified under the LMIC design. Genetic evaluation with phenotyped cows from intermediate and large average herd sizes had a higher accuracy than genetic evaluation with phenotyped cows from small average herd sizes. Increases in average herd size had a diminishing effect on increases in accuracy of genetic evaluation of phenotyped cows. The genetic evaluation of phenotyped cows using genomic information had higher accuracy compared to pedigree information across all breeding designs. Table 1 reports the accuracy of EBVs of phenotyped cows with both genetic evaluation methods as average herd size was changed. The accuracies reported correspond to models with the herd modelled as a random effect. At an average herd size of 1 (μ=1.58), phenotyped cows had an accuracy of EBVs of 0.40 with the PBLUP and 0.50 with the GBLUP (an increase of 0.10). At all other average herd sizes, the increase in accuracy of GBLUP compared to PBLUP was between 0.11 and 0.12. In what follows, results will only be presented for the GBLUP.

**Table 1.** The impact of genetic evaluation method on EBV accuracy.

| *Method* |  | *Size of Herd* |  |  |  |  |  |
| --- | --- | --- | --- | --- | --- | --- | --- |
|  |  | *1* | *2* | *4* | *8* | *16* | *32* |
| *PBLUP* | *Accuracy* | *0.40* | *0.41* | *0.43* | *0.44* | *0.45* | *0.46* |
| *GBLUP* | *Accuracy* | *0.50* | *0.53* | *0.54* | *0.56* | *0.57* | *0.57* |

*Comparison of the accuracy of genetic evaluation method under the LMIC design with different average herd sizes and using the PBLUP or GBLUP method. Herd is modelled as a random effect. Standard error was 0.01 or less.*

### Impact of herd effect modelling

Genetic evaluations were run using three models: (i) excluding a herd effect, (ii) herd modelled as a fixed effect, and (iii) herd modelled as a random effect. The genetic evaluation of phenotyped cows that included a herd effect had higher accuracies across all breeding designs. The genetic evaluation of phenotyped cows with genomic information and modelling herd as a random effect had higher accuracy compared to modelling herd as a fixed effect at low average herd sizes. However, the accuracies of the two modelling approaches converged once a herd size of 8 was reached. Figure 2 plots the average herd size against the accuracy for each of the three evaluation models. Figure 2 shows that excluding a herd effect gave an accuracy of 0.48, averaged across all herd sizes. At average herd sizes of 1.58 and 2.32, modelling herd as a random effect increased the accuracy by 0.10 and 0.05, compared to modelling herd as a fixed effect. At an average herd size of 8, the accuracies from the two modelling approaches had practically converged.

**Figure 2.**
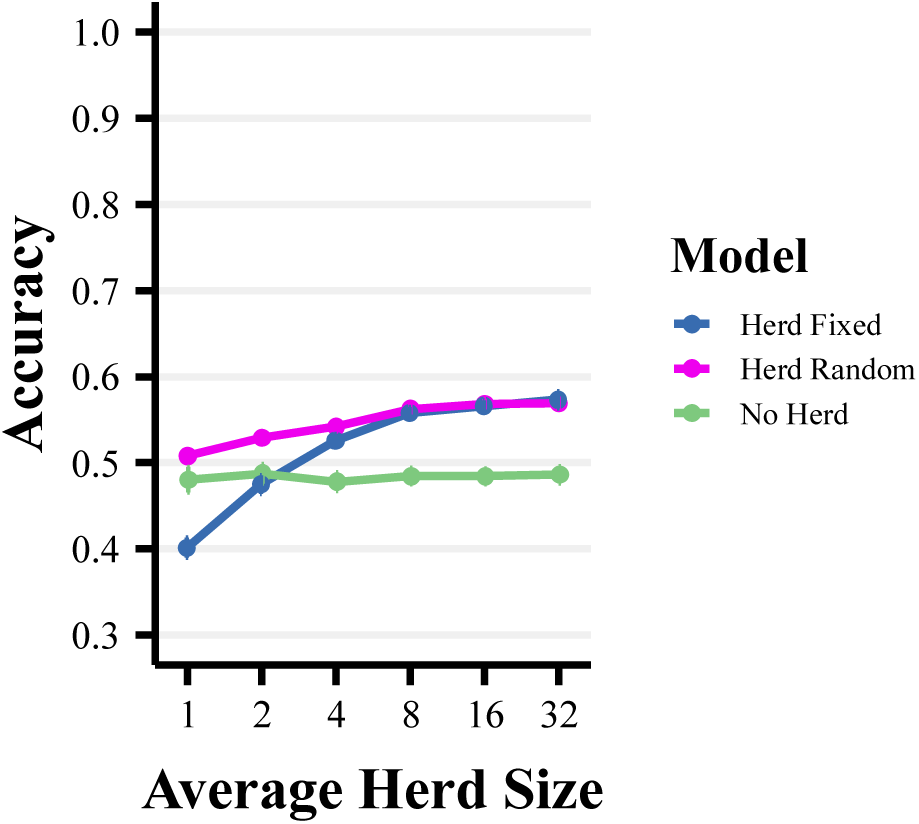
The impact of the model on EBV accuracy of cows. Comparison of the statistical modelling of herd under the LMIC design with GBLUP. The accuracy of estimated breeding values as a function of average herd size (1-32) and the herd effect (i) excluded from the model, (ii) modelled as a fixed effect and (iii) modelled as a random effect.

### Impact of genetic connectedness and trait heritability

In the simulations we varied genetic connectedness between herds in two ways; (i) herd connectivity – varied by simulating different average herd sizes; and (ii) the recent N_e_ of the breeding design - varied using different numbers of sires mated per generation. The genetic evaluation of phenotyped cows from breeding designs with strong genetic connectedness had higher accuracy compared to breeding designs with weaker genetic connectedness. Figure 3 plots the average herd size against the accuracy of EBVs of phenotyped cows for each of the four breeding designs with different numbers of sires mated per generation. Figure 3 shows that at an average herd size of 1 (μ=1.58), a decrease in the number of sires mated per generation from 5,000 to 1,000, 250 and 100 increased the accuracy from 0.46 to 0.50, 0.55 and 0.62, respectively. This shows the individual impact of the number of sires mated per generation on the accuracy. With 1,000 sires mated per generation, an increase in the average herd size from 1.58 to 32, increased the accuracy from 0.50 to 0.58. This shows the individual impact of the average herd size on the accuracy. An increase in the average herd size from 1.58 to 32, and a decrease in the number of sires mated per generation from 1,000 to 100, increased the accuracy from 0.50 to 0.68. This shows the combined impact of the genetic connectedness of the breeding design on the accuracy.

**Figure 3.**
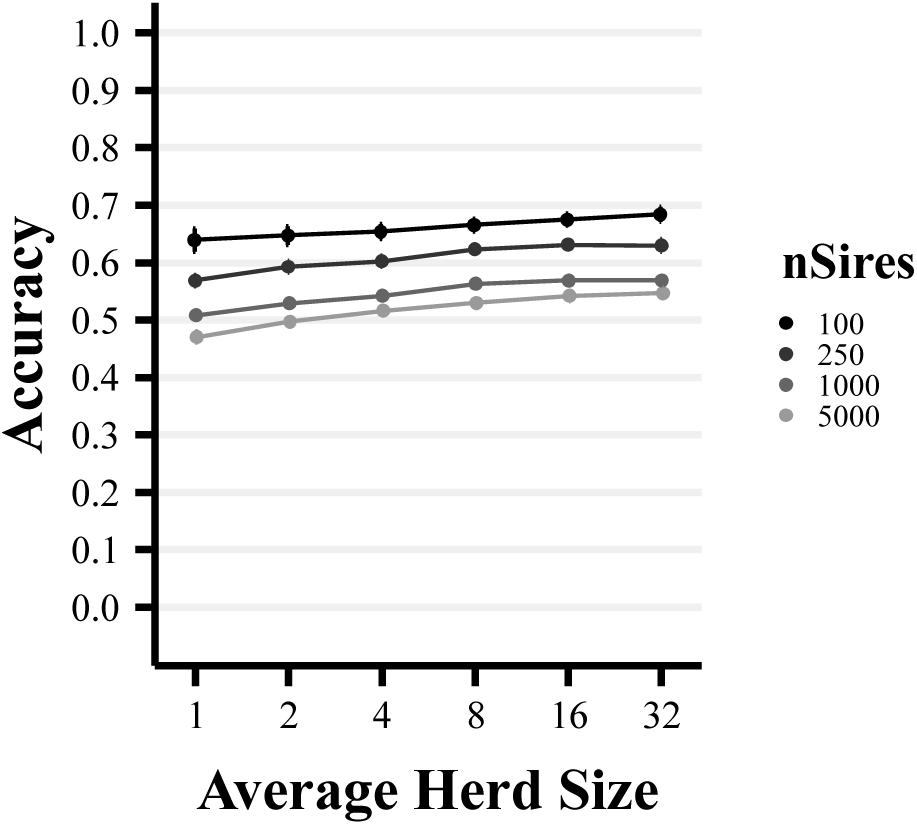
The impact of genetic connectedness on EBV accuracy of cows. Comparison of genetic connectedness of the training set with GBLUP. The accuracy of estimated breeding values are presented as a function of average herd size (1-32) and the number of sires (100, 250, 1000 & 5000). The number of records in the training set is 8000. Herd is modelled as a random effect.

The genetic connectedness of the breeding design also showed interactions with the heritability of the trait. Across all trait heritabilities, the EBVs of phenotyped cows had lower accuracy in breeding designs that had weak genetic connections. The lower accuracy due to an increase in the number of sires mated per generation in the breeding design became more prominent at lower heritabilities. The lower accuracy due to a decrease in the average herd size of the breeding design was more prominent at higher heritabilities. Figure 4 plots the average herd size against the accuracy of EBVs of phenotyped cows for two of the four different numbers of sire mated per generation (100 and 1,000 sires). The three panels correspond to the heritability under the different breeding designs. Figure 4 shows that the highest accuracy (0.94) was achieved for a high heritability trait (0.5) and when genetic connectedness was strong (100 sires mated per generation and an average herd size of 32). A decrease in the average herd size from 32 to 1.58, reduced the accuracy by 0.07. An accuracy of 0.68 was achieved for a low heritability trait (0.1) and when genetic connectedness was strong (100 sires mated per generation and an average herd size of 32). An increase in the number of sires mated per generation to 1,000 sires mated per generation, reduced the accuracy by 0.10.

**Figure 4.**
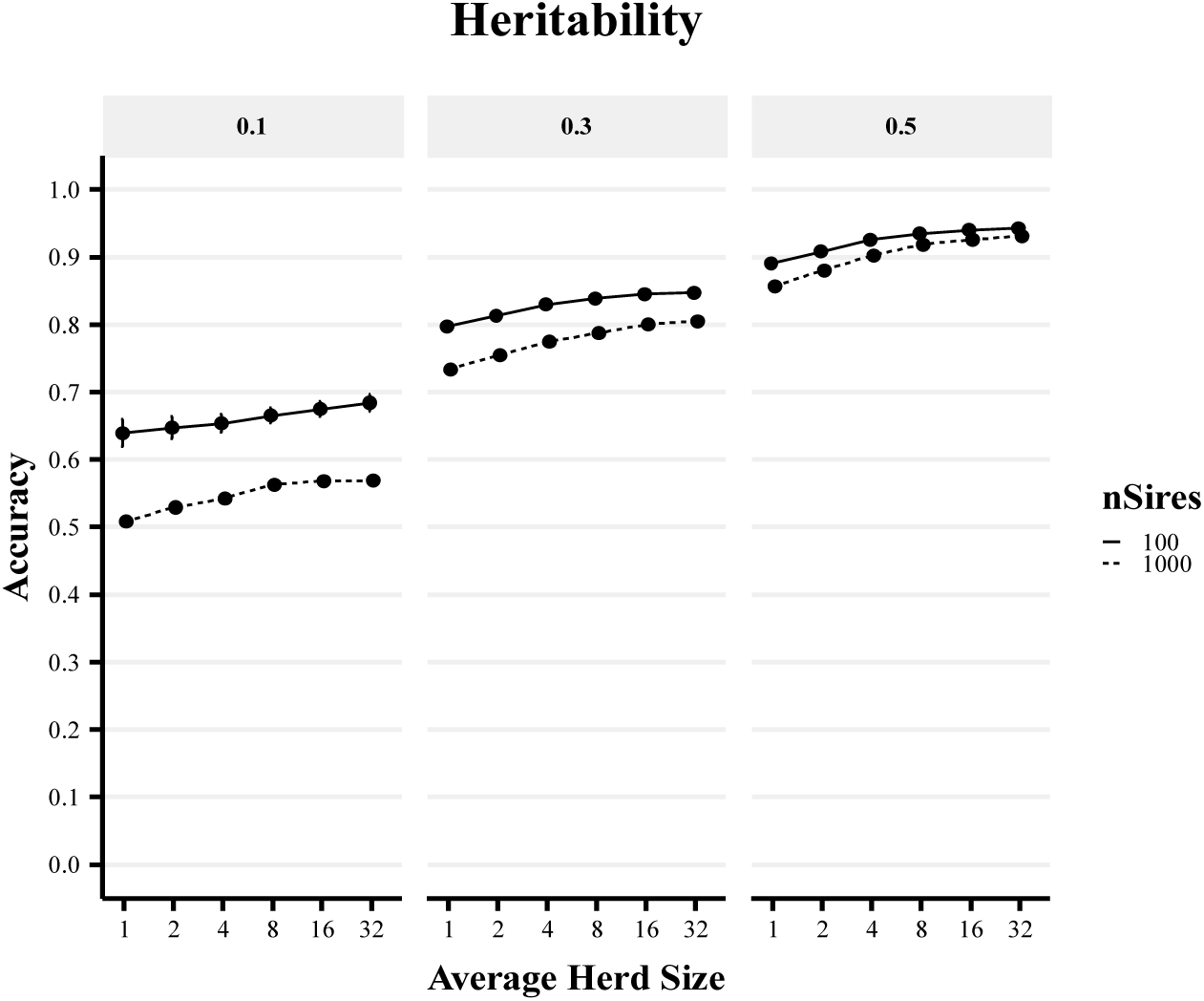
The impact of genetic connectedness and heritability on EBV accuracy of cows. Comparison of the heritability of the trait and genetic connectedness with GBLUP. The accuracy of estimated breeding values as a function of average herd size (1-32) and the genetic connectedness of the training set (100 & 1,000 sires per generation). The three panels correspond to the heritability of the trait (0.1, 0.3 & 0.5). Herd is modelled as a random effect.

### Impact of Training Set Size

Genetic evaluation of phenotyped cows with a larger number of records had higher accuracies for all average herd sizes. Figure 5 plots the average herd size against the accuracy of EBVs of phenotyped cows for the four different training set sizes. Figure 5 shows an increase in the number of records in the training set increased the accuracy across all of the average herd sizes. At an average herd size of 1 (μ=1.58), an increase in the number of records in the training set from 2,000 to 8,000, 16,000 and 32,000 records increased the accuracy from 0.41 to 0.50, 0.59 and 0.68, respectively.

**Figure 5.**
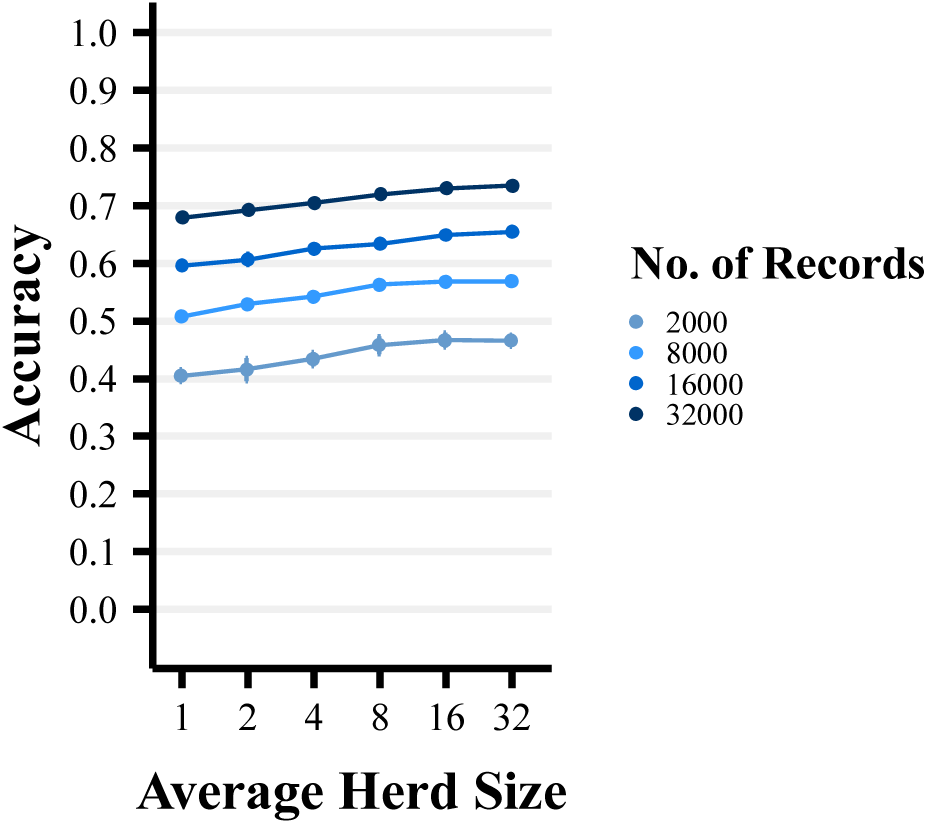
The impact of training set size on EBV accuracy of cows. Comparison of the number of records in the training set with GBLUP. The accuracy of estimated breeding values of cows as a function of average herd size (1-32) and the number of records in the training set (2000, 8000, 16000 & 32000). Herd is modelled as a random effect.

### Prediction of young bulls

Genomic prediction of young bulls was possible using marker estimates from the genetic evaluations of their phenotyped dams. The accuracies of young bulls followed similar trends to those observed in the evaluation of phenotyped cows, with a reduction of ∼0.1 in overall accuracy. Genomic prediction of young bulls with a larger number of records in the training set had higher accuracies. The accuracy of genomic prediction of young bulls from an average herd size of 1 (μ=1.58) was 0.40 under a breeding design with 1,000 sires mated per generation and a training set of 8,000 phenotyped and genotyped cows. Figure 6 plots the accuracy of EBVs of candidate young bulls against the average herd size for the four different training set sizes. Figure 6 shows that an increase in the number of records in the training set increased the accuracy across all of the average herd sizes. At an average herd size of 1 (μ=1.58), an increase in the number of records in the training set from 2,000 to 8,000, 16,000 and 32,000 records increased the accuracy from 0.28 to 0.40, 0.51 and 0.62, respectively.

**Figure 6.**
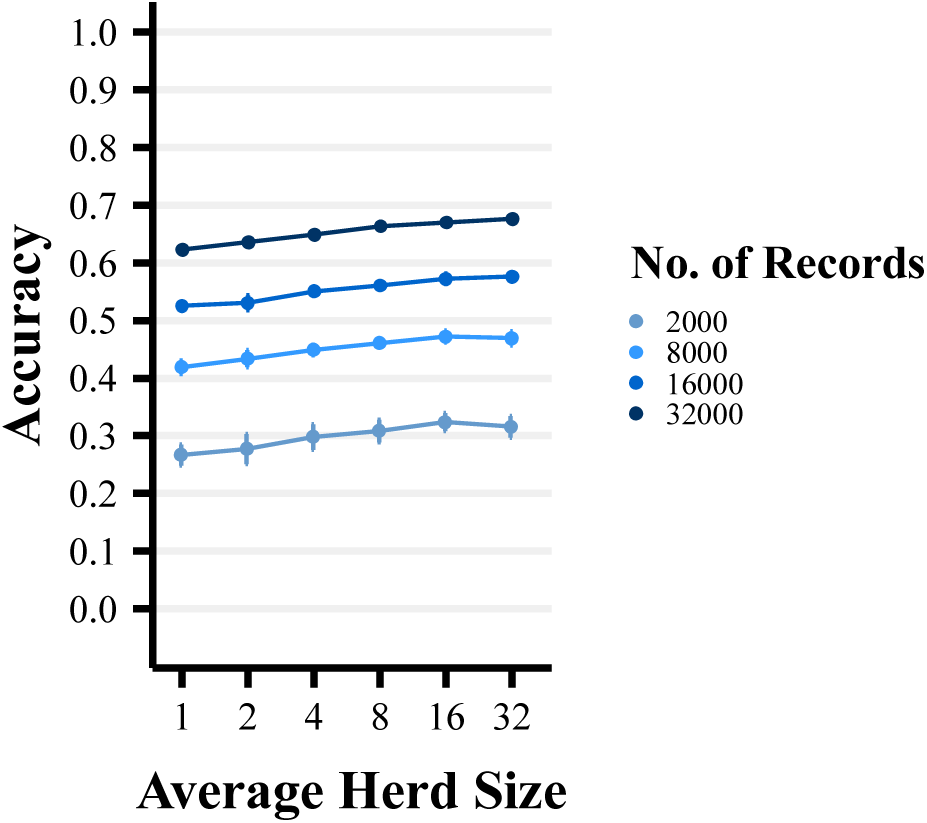
The impact of training set size on EBV accuracy of young bulls. Comparison of the number of records in the training set and genetic connectedness with GBLUP. The accuracy of genomic estimated breeding values of young bulls as a function of average herd size (1-32) and the number of records in the training set (2000, 8000, 16000 & 32000. Herd is modelled as a random effect.

The accuracy was also affected by an interaction between the heritability of the trait and the genetic connectedness of the breeding design. The genetic connectedness of the breeding design was less important for traits with a higher heritability. Figure 7 plots the accuracy against the average herd size for two of the four different numbers of sire mated per generation (100 and 1,000 sires). The three panels correspond to the different trait heritabilities in the breeding designs. Figure 7 shows that an increase in the average herd size did not recover the loss of accuracy due to lower genetic connectedness (100 vs 1,000 sires mated per generation). This is different from what was observed with the accuracy for phenotyped cows. Figure 7 shows that for a high heritability trait (0.5) and an average herd size of 32, increasing the number of sires mated per generation from 100 to 1,000 sires mated per generation reduced the accuracy of young bulls by 0.04.

**Figure 7.**
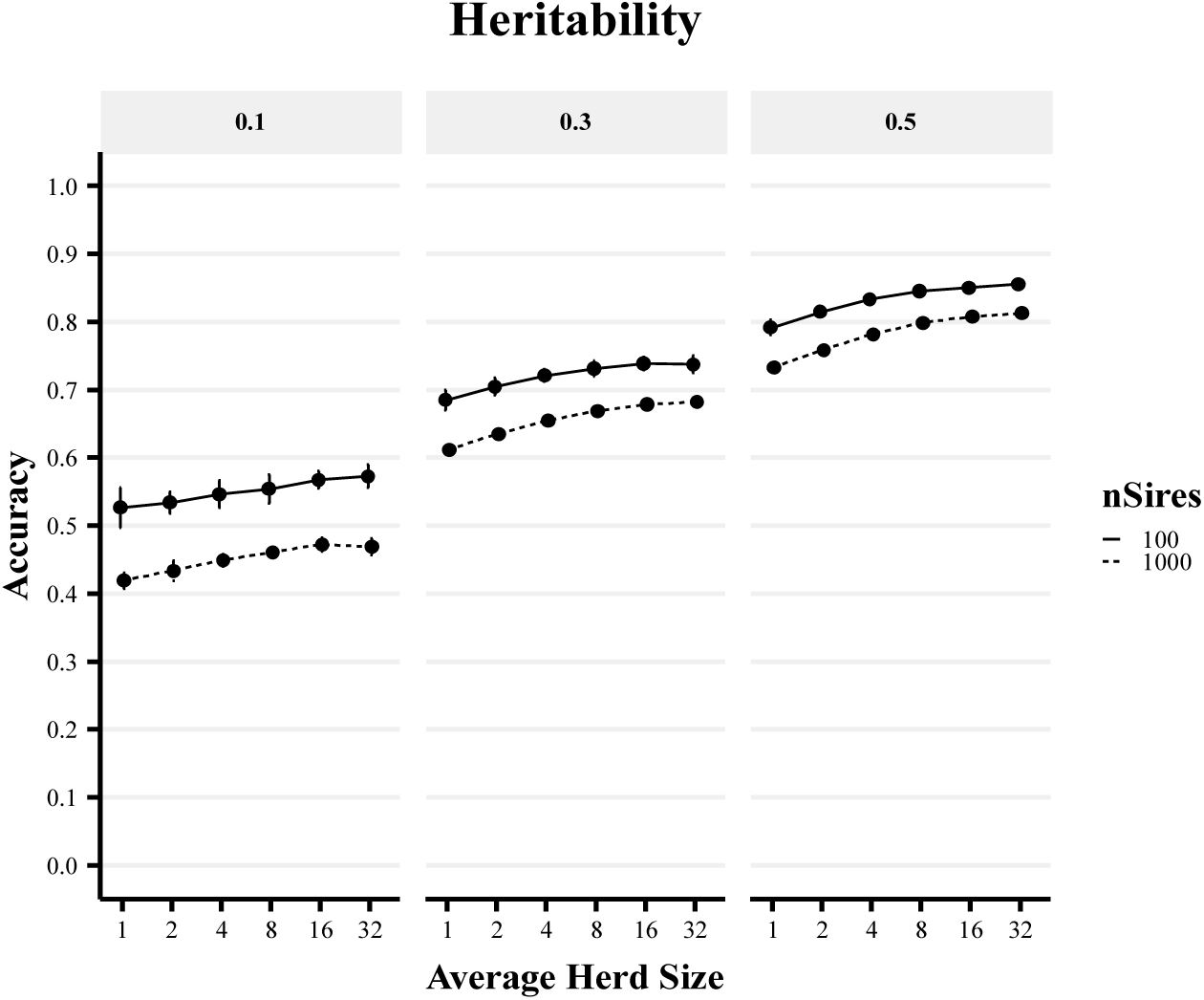
The impact of genetic connectedness and heritability on EBV accuracy of young bulls. Comparison of the heritability of the trait with GBLUP. The accuracy of genomic estimated breeding values of young as a function of average herd size (1-32) and the genetic connectedness of the training set (100 & 1,000 sires per generation). The three panels correspond to the heritability of the trait (0.1, 0.3 & 0.5). Herd is modelled as a random effect.

## Discussion

In this paper, we demonstrated that genetic evaluation using genomic information can provide accurate EBVs when using data recorded on smallholder farms across a range of breeding designs. Therefore, genetic evaluations using genomic information could enable *in-situ* data recorded on smallholder farms to be used to drive *in-situ* genetic improvement programs and genetic importation programs to improve animal performance on such smallholder farms. This capacity would enable tailored improvement and importation of genetics for smallholder farms. The results of our study highlight three main points for discussion: (i) factors that impact the accuracy of genomic evaluations; (ii) limitations of the simulation; and (iii) prospects for animal breeding in LMIC smallholder dairy production systems.

### Factors that impact the accuracy of genomic evaluations

#### Impact of Herd Size

The herd or management group is usually included in the statistical model of genetic evaluations to enhance the partitioning of the genetic merit of an individual from the non-genetic effects underlying its phenotype [21–24]. Herds can be modelled as fixed or random effects. One of the reasons underlying the great success of genetic evaluations in advanced economies is that large data sets are routinely assembled from commercial farms with large herd sizes. This data structure is suited to modelling herd as a fixed effect. This data structure also enables accurate separation of genetic and environmental effects and reduces potential bias due to a difference in management effects between different herds.

However, LMIC smallholder dairy farms often have small herd sizes, typically between one and five cows. With herd sizes as small as this, LMIC smallholder dairy datasets sit at one extreme of the bias-variance trade-off [34]. Modelling herd as a fixed effect provides unbiased estimates. However, when herd sizes are small, these estimates of herd effect may have large variance. Therefore, modelling herd as a fixed effect in the LMIC smallholder dairy genetic evaluations may lead to herd effect estimates with high variance and a reduced ability to correctly rank individuals by genetic merit [25]. This could lead to a decreased accuracy of EBVs. An alternative approach in such settings would be to model herds as random effects. Modelling herd as a random effect looks to minimize the variance of estimates, but the resulting estimates are inherently biased due to shrinkage applied during estimation. However, the shrinkage process allows phenotypes recorded in small herds to partially and proportionately contribute to the genetic evaluation. This is essential for LMIC smallholder dairy genetic evaluations with herd sizes typically between one and five cows. The results from our study support this and showed that when data is collected from herds between one and four cows, genomic evaluations modelling herd as a random effect outperformed modelling herd as a fixed effect. In the case of genomic evaluations using data from an average herd size of 1 (μ=1.58), modelling herd as a random effect increased the accuracy of EBVs of phenotyped cows by 0.10 compared to modelling herd as a fixed effect. It was only when the average herd size was 8 or more that the accuracy of EBVs of phenotyped cows from the two models converged. Overall our results demonstrate that modelling herd as a random effect in LMIC smallholder dairy genetic evaluations: (i) increases the accuracy of genetic evaluations; (ii) enables phenotypes recorded in all herds to partially and proportionately contribute to the genetic evaluation; and (iii) enables the breeding values of all animals (even those in single cow herds) to be calculated. However, as is discussed later, modelling herd as a random effect may increase accuracy but bias may be generated when non-random associations between the genetic value of cattle and the herd management exist within the training set.

#### Impact of GBLUP as a tool to increase connectedness between herds

Sufficient genetic connectedness between herds is important for accurate genetic evaluations [16,35]. In dairy production systems in advanced economies, large herd sizes and widespread use of artificial insemination creates strong genetic connectedness between herds that enables accurate separation of genetic and environmental effects. Because strong genetic connectedness between herds is already established in dairy production systems in advanced economies, GBLUP has primarily increased the accuracy of EBVs compared to PBLUP by capturing and exploiting deviations from expected relationships between cattle caused by Mendelian sampling [36–38]. For example, the accuracy, which is the square root of reliability, of prediction for milk yield of young bulls have increased from 0.62 using PBLUP to 0.85 for GBLUP [20]. We say “primarily” because most training populations are comprised of bulls that were progeny tested across a large number of herds. In this situation, modelling both the genetic and herd effects jointly is less of a concern. The single-step GBLUP method and the recent rise of cow genotyping will also enable improvements by jointly modelling of genetic and herd effects. In LMIC smallholder dairy production systems the benefit using GBLUP will be both due to exploiting deviations from expected relationships caused by Mendelian sampling and due to implicit increases of genetic connectedness between herds.

Generating sufficient genetic connectedness between herds is especially difficult and important in LMIC smallholder dairy production systems because herd sizes are often small, farms are geographically dispersed, and artificial insemination is not widely used [8]. In such production systems, the genetic and environmental effects are likely to be partially or fully confounded. This is most obvious in the case of a single cow herd where we cannot separate the genetic effect of the cow from the herd effect of the farm. However, a range of levels of confounding could also arise in small herds composed of cows sharing the same pedigree-derived relatedness, with the recent common ancestor or ancestors only used in that herd. In both of these circumstances, PBLUP has limited ability to partition a cow’s phenotype into its genetic and environmental components. In contrast, GBLUP can achieve this partitioning, because it is capable of tracking the different permutations of haplotypes shared between cattle in different herds. During a genetic evaluation, GBLUP implicitly estimates the effects of these haplotypes and from this also the EBV of each animal. This allows phenotypic records from cows with shared haplotypes in different herds to contribute to the implicit estimation of haplotype effects and the estimates of those haplotype effects allows the partitioning of those cow’s phenotypes into their genetic and herd environment components. Furthermore, through this implicit increasing of genetic connectedness between herds, GBLUP increases the number of herds and cows that contribute useable information to the genetic evaluation compared to PBLUP. All of these interlinked factors that underlie the advantages of GBLUP, firstly make genetic evaluations using data recorded *in-situ* on smallholder herds possible, and secondly, work to make those genetic evaluations more accurate than those of PBLUP. In our study, the increase in genetic connectedness provided by GBLUP resulted in genetic evaluations with approximately 0.1 higher accuracy of EBVs compared to PBLUP, independent of herd size. This result probably overestimates the power of PBLUP in such settings. We used five generations of error-free pedigree records in PBLUP. In reality, limited pedigree recording takes place in LMIC smallholder dairy production systems. We should emphasise though that LMIC smallholder dairy data structures likely do not enable very accurate estimation of individual haplotype effects and that the dataset size will continue to be an important factor.

Another benefit of the increased genetic connectedness of training sets provided by GBLUP, not assessed in our study, may be the mitigation of the bias of EBVs. In LMIC smallholder dairy production systems, natural sire mating is prevalent, pedigree recording is limited, herd sizes are often small and farms are geographically dispersed. This structure is likely to lead to isolated family clusters in pedigrees. Therefore, when using PBLUP in LMIC smallholder dairy genetic evaluations, most of the information used to calculate the EBV for any particular individual will be provided by close relatives captured by this poorly connected pedigree. This may result in only a very small number of herds contributing effective information to the genetic evaluation of an animal or group of related animals. This becomes a problem if confounding exists between the environment and the genetics in the isolated clusters of herds. Confounding can occur when the same natural service bull is used by a cohort of farmers with farms that have a better or worse than average herd environment. This may lead to biased breeding values under PBLUP. In contrast, haplotypes are likely to be dispersed across more herds. Therefore, GBLUP could accumulate effective information from more herds and more cows and thus be less prone to having haplotypes confounded with the environment.

### Limitations of the simulation

Our simulations did not model the full complexity that would arise in practical genetic evaluations for LMIC smallholder dairy production systems. In this section we discuss three limitations of our simulations: (i) high genomic selection accuracy; (ii) a simplified distribution of animals across farms; and (iii) a simplified breeding goal.

#### Impact of high genomic selection accuracy

The accuracies of EBVs of phenotyped cows and young bulls observed in these simulations are likely higher than what may be expected in practical genetic evaluations for LMIC smallholder dairy production systems. Several simplifications of the simulation are likely to have caused this, including the absence of genotyping and pedigree errors, additive genetic architecture, homogeneity of environment and a single breed. Also, fixed variance components were used in the estimation of EBVs. In practical LMIC genetic evaluations, the estimation error of variance components may result in lower accuracies of EBVs. However, we believe that the main conclusion from this study (i.e., that GBLUP is more powerful than PBLUP in LMIC smallholder production systems for several reasons) would still hold for more realistic simulations or real data. For decades it has been difficult to sustain widespread recording and use of pedigree to drive genetic evaluations in LMIC dairy production systems. GBLUP, for the reasons we outline, offers a route to overcoming this problem.

#### Impact of simplified distribution of animals across farms

The distribution of cattle across herds in the population impacts the choice of modelling herd as a fixed or random effect in genetic evaluations. Bias, detected in this study as an inflation or deflation of EBVs, can be generated when a non-random association between herd management and genetic potential of cattle exists. Such non-random associations can be generated, for example, by well-resourced farmers who use better management practices also being able to afford semen of higher genetic merit sires, or by the restriction of natural mating sires to herds in specific regions. As discussed previously, modelling herd as a fixed effect estimates the herd effects independently for each herd. When herd sizes are large, such as in advanced economies, this can reduce bias caused by differences in the genetic means of different herds. Herd sizes are not large in LMIC smallholder dairy production systems. In such circumstances, modelling herd as a random effect in genetic evaluations allows phenotypes recorded in small herds to partially and proportionately contribute to the genetic evaluation. This benefit extends to small herds composed of cows of varying relatedness, with the ancestral haplotypes only present in that herd. This is important in an LMIC smallholder dairy production systems context, with more than 70% of milk in Kenya produced by herds of one to five cows [6,7]. However, the choice between modelling herd as a random effect should consider the bias-variance trade-off [34].

This choice is particularly important if correlations between herd management and the genetic value of cows exist. Under this scenario, if the differences in genetic means across herds are not accounted for, the herd effect of an animal may be partially assigned to the genetic effect when herd is modelled as a random effect. In our study, cattle were assigned to herds at random and no correlation between herd management and the genetic value of cows existed. Therefore, significant bias effects were only detected in genetic evaluations modelling herd as a fixed effect with an average herd size of one (results not shown). There is another impact of the simulation not modelling the full complexity of the distribution of cattle and its genetic effects across farms. The training sets likely had an increased genetic connectedness compared to practical genetic evaluations in LMIC smallholder dairy production systems. This resulted in accuracies of EBVs that are likely to be higher than expected in practical genetic evaluations in LMIC smallholder dairy production systems. However, our study also did not capture the full complexity of the interaction between genetic connectedness and herd size. Therefore, our results likely underestimated the benefits of GBLUP to increase genetic connectedness and more accurately separate the genetic and environmental components of each cow’s phenotype in small herds in practical genetic evaluations in LMIC smallholder dairy production systems. With the projected increases in data recording, we expect that these effects will diminish or that the scale of the data will enable at least reasonably high accuracy to stimulate genetic progress.

#### Impact of simplified breeding goal

The breeding program examined in this simulation only considered a single quantitative trait that did not interact with the environment. The breeding goal for practical LMIC smallholder dairy production systems would be much more complex in practice. It would comprise of several correlated traits (e.g., milk yield, milk components, fertility, feed requirements, heat tolerance, disease resistance) many of which would interact with the environment. The single quantitative trait with 10,000 QTL that we simulated is representative of such an index with a few additional assumptions: all traits are measured on all animals, all traits are pleiotropic, and economic merit is linear. This study simulated a simplified genetic architecture without considering dominance, epistasis and gene by environment interaction. This will likely decrease the absolute values of accuracy reported in this study but the main conclusions of our study (i.e., that GBLUP is more powerful than PBLUP in LMIC smallholder dairy production systems for several reasons) will still hold.

### Prospects for animal breeding in LMICs

Our motivations for undertaking this study were to contribute to the enabling of the sustained and long-term use of animal breeding to improve agricultural productivity and sustainability in LMIC smallholder dairy production systems. Breeding has been hugely successful for improving animals and plants in advanced economies and for improving plants in LMICs. Breeding has had limited success in improving animals in LMICs. We believe that for animal breeding to be successful in LMIC smallholder dairy production systems it must be driven by data recorded *in-situ* on animals from such farms. We believe that the limited success of animal breeding in these contexts is due to the infrastructure and data structures that are prevalent in these systems, which make genetic evaluation using pedigree difficult, if not impossible. Specifically, the infrastructure required to record pedigree over long periods of time is typically absent in LMIC smallholder dairy production systems. The lack of widespread use of AI and the small herd sizes result in a data structure that has insufficient genetic connectedness between herds to facilitate genetic evaluations based on pedigree. We believe that genomic data offers a route to overcome these problems and the results of our study show this. However, our study did not quantify the long-term impacts of genomic data in LMIC smallholder dairy breeding programs. As an example, our study demonstrated that the EBVs of young bulls from an average herd size of 1 (μ=1.58) could be predicted with an accuracy of 0.40. However, as well as increasing the accuracy of selection, genomic evaluations also offer an opportunity to reduce the generation interval of breeding programs. These reductions in the generation interval have been the primary driver of the gain in the rate of genetic improvement in dairy breeding programs in advanced economies because they have approximately halved the generation interval, thereby doubling the rate of genetic gain [20]. In LMIC breeding programs, it is difficult to estimate the reductions in the generation interval that genomic evaluations could provide. This is due to the lack of pedigree recording and infrastructure for the widespread use of AI, already discussed. However, it is possible to say that genomic evaluations will allow LMIC breeding programs to drive the generation interval to near the biological and economic minimum for that system. The impact of this, and the other results from our study, on the long-term genetic gain of LMIC smallholder dairy breeding programs will need to be explored further.

Genomic data is expensive and its requirement may create a new cost barrier to the success of animal breeding in LMIC smallholder dairy production systems. New business models are needed to overcome this barrier in a self-sustaining way. One such model could involve establishing an intertwined breeding and dissemination program for a target environment. The cost of operating the breeding program would need to be proportionate to the market that it would serve via its dissemination program. The breeding program could comprise an informal set of nucleus animals distributed across many small herds within the target environment. These nucleus animals could be genotyped and phenotyped and this data used for a genetic evaluation using GBLUP. The best animals from this nucleus could be disseminated via artificial insemination (with or without a subsequent progeny testing scheme), as natural service sires, or as heifers. Further, the genomic prediction equation calculated for the genetic evaluation could be used to select any external animals that would be imported into the region. To reduce the costs of data recording in the nucleus and to increase the value of what would be disseminated a whole range of additional technologies and services could be bundled together. For example, nucleus herds could also serve as demonstration herds and the dissemination program could provide additional extension services (e.g., a text message for a small fee with management or market information). Or improved animal genetics could be packaged together with other technology (e.g., improved seeds) which may have higher adoption rates. Overall, a business model could be constructed that bundles technology, data recording, extension services, and a marketplace for LMIC smallholder farmers. This type of self-sustaining platform would maximize the benefits and cost-efficiency of any component (e.g., the genotyping and phenotyping of animals). This business model could leverage the successes of established technologies and practices to drive adoption of those that have been traditionally more intractable. The Africa Dairy Genetic Gains [14], the Public Private Partnership for AI Dissemination [15] projects and the emerging social enterprises (e.g., One Acre Fund [39], and electronic marketplaces for agricultural products in LMICs (e.g., Livestock 247 [40]) show that many components of such a model are already in place.

## Conclusions

This study has demonstrated the potential of genomic information to be an enabling technology in LMIC smallholder dairy production systems by facilitating genetic evaluations with *in-situ* records collected from farms with herd sizes of four cows or less. Across a range of breeding designs, genomic data made it possible to accurately predict EBVs of phenotyped cows and young bulls using data sets that contained small herds that had weak genetic connections. The use of *in-situ* smallholder dairy data in genetic evaluations would establish breeding programs to improve *in-situ* germplasm and, if required, would enable the importation of the most suitable external germplasm. This could be individually tailored for each target environment. Together this would increase the productivity, profitability and sustainability of LMIC smallholder dairy systems. However, genomic data is expensive and business models will need to be carefully constructed so that the costs are sustainably offset.

## Declarations

### Ethics approval and consent to participate

Not applicable.

### Consent for publication

Not applicable.

### Availability of data and material

The simulated data and materials are availailable upon request.

### Author’s contributions

JMH conceived the study. JMH and OP designed the study. OP performed the analysis. OP and JMH wrote the manuscript. RM, RCG, MJ and GG helped interpret the result and refined the manuscript. All authors read and approved the final manuscript.

## Acknowledgements

The authors thank Georgios Banos for valuable comments on a previous draft of the paper. This work has made use of the resources provided by the Edinburgh Compute and Data Facility (ECDF) (http://www.ecdf.ed.ac.uk).

## References

1. Dekkers J, Hospital F. The use of molecular genetics in the improvement of agricultural populations. Nat Rev Genet 2002;162:1945–59. doi:10.1038/nrg701.

2. Kahi AK, Nitter G, Gall CF. Developing breeding schemes for pasture based dairy production systems in Kenya: II. Evaluation of alternative objectives and schemes using a two-tier open nucleus and young bull system. Livest Prod Sci 2004;88:179–92. doi:10.1016/j.livprodsci.2003.07.015.

3. Muriuki HG. Dairy development in kenya. Food Agric Organ United Nations 2011:1– 52.

4. Ojango JMK, Wasike CB, Enahoro DK, Okeyo AM. Dairy production systems and the adoption of genetic and breeding technologies in Tanzania, Kenya, India and Nicaragua. Anim Genet Resour 2016;59:81–95. doi:10.1017/S2078633616000096.

5. FAOSTAT. FAOSTAT 2018. http://www.fao.org/faostat/ (accessed November 1, 2018).

6. East African Dairy Development Program. The Dairy Value Chain in Kenya 2012:2011–2.

7. Abdulsamad A, Gereffi G. East Africa dairy value chains: Firm capabilities to expand regional trade. 2016.

8. Ojango JMK, Marete A, Mujibi D, Rao J, Pool J, Rege JEO, et al. A novel use of high density SNP assays to optimize choice of different crossbred dairy cattle genotypes in small-holder systems in East Africa. Proc 10th World Congr Genet Appl to Livest Prod 2014:2–4. doi:10.13140/2.1.3730.5603.

9. Mutavi SK, Kanui TI, Njarui DM, Musimba NR, Amwata DA. Innovativeness and Adaptations : The Way forward for Small scale Peri-Urban Dairy Farmers in Semi-Arid Regions of South Eastern Kenya 2016;3:1–14.

10. Ducrocq V, Laloe D, Swaminathan M, Rognon X, Tixier-Boichard M, Zerjal T. Genomics for ruminants in developing countries: From principles to practice. Front Genet 2018;9:1–7. doi:10.3389/fgene.2018.00251.

11. Rothschild MF, Plastow GS. Applications of genomics to improve livestock in the developing world. Livest Sci 2014;166:76–83. doi:10.1016/j.livsci.2014.03.020.

12. Potdar V, Bhave K, Khadse J. BAIF Experience in Field Data Collection International Journal of Animal 2017:3–7.

13. Wurzinger M, Sölkner J, Iñiguez L. Important aspects and limitations in considering community-based breeding programs for low-input smallholder livestock systems. Small Rumin Res 2011;98:170–5. doi:10.1016/j.smallrumres.2011.03.035.

14. ADGG. Africa Dairy Genetic Gains (ADGG) n.d. https://africadgg.wordpress.com/.

15. PAID. Public Private Partnership n.d. https://www.slideshare.net/ILRI/adgg-paid2-feb2017 (accessed August 14, 2019).

16. Kennedy BW, Trus D. Considerations on genetic connectedness between management units under an animal model. J Anim Sci 1993;71:2341–52. doi:10.2527/1993.7192341x.

17. Mrode R, Ojango JMK, Okeyo AM, Mwacharo JM. Genomic Selection and Use of Molecular Tools in Breeding Programs for Indigenous and Crossbred Cattle in Developing Countries: Current Status and Future Prospects. Front Genet 2019;9. doi:10.3389/fgene.2018.00694.

18. Nejati-Javaremi a., Smith C, Gibson JP. Effect of total Allelic Relationship on Accuracy of Evaluation and Response to Selection. J Anim Sci 1997;75:1738–45. doi:10.1111/j.1540-5834.2011.00603.x.

19. Villanueva B, Pong-Wong R, Fernandez J, Toro MA. Benefits from marker-assisted selection under an additive. J Anim Sci 2005;83:1747–52.

20. Wiggans GR, Cole JB, Hubbard SM, Sonstegard TS. Genomic Selection in Dairy Cattle: The USDA Experience. Annu Rev Anim Biosci 2017;5:309–27. doi:10.1146/annurev-animal-021815-111422.

21. Schaeffer LR. Contemporary Groups Are Always Random. Personal 2009. http://www.aps.uoguelph.ca/~lrs/LRSsite/ranfix.pdf (accessed August 14, 2019).

22. Visscher MP., Goddard ME. Fixed and Random Contemporary Groups. J Dairy Sci 1993;76:1444–54.

23. Ugarte E, Alenda R, Carabaño MJ. Fixed or Random Contemporary Groups in Genetic Evaluations. J Dairy Sci 1992;75:269–78. doi:10.3168/jds.S0022-0302(92)77762-5.

24. Frey M, Hofer A, Künzi N. Comparison of models with a fixed or a random contemporary group effect for the genetic evaluation for litter size in pigs. Livest Prod Sci 1997;48:135–41.

25. Oikawa T, Sato K. Treating small herds as fixed or random in an animal model. J Anim Breed Genet 1997;114:177–83. doi:10.1111/j.1439-0388.1997.tb00503.x.

26. Gaynor RC, Gorjanc G, Wilson D, Money D, Hickey JM. AlphaSimR 2019.

27. Chen GK, Marjoram P, Wall JD. Fast and flexible simulation of DNA sequence data. Genome Res 2009;19:136–42. doi:10.1101/gr.083634.108.

28. MacLeod IM, Larkin DM, Lewin HA, Hayes BJ, Goddard ME. Inferring demography from runs of homozygosity in whole-genome sequence, with correction for sequence errors. Mol Biol Evol 2013;30:2209–23. doi:10.1093/molbev/mst125.

29. Gibson J, Ojango J, Okeyo M. Dairy Genetics East Africa (DGEA) Phase 2 - The final narrative report to the Bill and Melinda Gates Foundation. 2014.

30. Henderson CR. Best Linear Unbiased Estimation and Prediction under a Selection Model. Biometrics 1975;31:423–47.

31. VanRaden PM. Efficient Methods to Compute Genomic Predictions. J Dairy Sci 2008;91:4414–23. doi:10.3168/jds.2007-0980.

32. Meyer K. WOMBAT – A tool for mixed model analyses in quantitative genetics by REML. J Zhejiang Uni Sci B 2007:815–21. doi:[doi:10.1631/jzus.2007.B0815].

33. Hickey JM, Gorjanc G. AlphaBayes n.d.

34. Stanek EJ, Well A, Ockene I. Why not routinely use best linear unbiased predictors (BLUPs) as estimates of cholesterol, per cent fat from kcal and physical activity? Stat Med 1999;18:2943–59. doi:10.1002/(SICI)1097-0258(19991115)18:21<2943::AID-SIM241>3.0.CO;2-0.

35. Foulley JL, Bouix J, Goffinet B, Elsen JM. Connectedness in Genetic Evaluation. In: Gianola D, Hammond K, editors. Adv. Stat. Methods Genet. Improv. Livest., Berlin, Heidelberg: Springer Berlin Heidelberg; 1990, p. 277–308. doi:10.1007/978-3-642-74487-7_13.

36. Hayes BJ, Visscher PM, Goddard ME. Increased accuracy of artificial selection by using the realized relationship matrix. Genet Res (Camb) 2009;91:47–60. doi:10.1017/S0016672308009981.

37. VanRaden PM, Van Tassell CP, Wiggans GR, Sonstegard TS, Schnabel RD, Taylor JF, et al. Invited Review: Reliability of genomic predictions for North American Holstein bulls. J Dairy Sci 2009;92:16–24. doi:10.3168/jds.2008-1514.

38. Su G, Guldbrandtsen B, Gregersen VR, Lund MS. Preliminary investigation on reliability of genomic estimated breeding values in the Danish Holstein population. J Dairy Sci 2010;93:1175–83. doi:10.3168/jds.2009-2192.

39. One Acre Fund n.d. https://oneacrefund.org/ (accessed August 14, 2019).

40. Livestock 247 n.d. https://livestock247.com/ (accessed August 14, 2019).

